# Autophagy is required to prevent early pro-inflammatory responses and neutrophil recruitment during *Mycobacterium tuberculosis* infection without affecting pathogen replication in macrophages

**DOI:** 10.1101/2022.11.04.515221

**Authors:** Rachel L. Kinsella, Jacqueline M. Kimmey, Asya Smirnov, Reilly Woodson, Margaret R. Gaggioli, Sthefany M. Chavez, Darren Kreamalmeyer, Christina L. Stallings

**Author notes:** Current address: Department of Microbiology and Environmental Toxicology, UC Santa Cruz, Santa Cruz, CA, USA. Correspondence: Rachel L. Kinsella, Christina L. Stallings.

## Abstract

The immune response to *Mycobacterium tuberculosis* infection determines tuberculosis disease outcomes, yet we have an incomplete understanding of what immune factors contribute to a protective immune response. Neutrophilic inflammation has been associated with poor disease prognosis in humans and in animal models during *M. tuberculosis* infection and, therefore, must be tightly regulated. ATG5 is an essential autophagy protein that is required in innate immune cells to control neutrophil-dominated inflammation and promote survival during *M. tuberculosis* infection, however, the mechanistic basis for how ATG5 regulates neutrophil recruitment is unknown. To interrogate what innate immune cells require ATG5 to control neutrophil recruitment during *M. tuberculosis* infection, we used different mouse strains that conditionally delete *Atg5* in specific cell types. We found that ATG5 is required in CD11c^+^ cells (lung macrophages and dendritic cells) to control the production of proinflammatory cytokines and chemokines during *M. tuberculosis* infection, which would otherwise promote neutrophil recruitment. This role for ATG5 is autophagy-dependent, but independent of mitophagy, LC3-associated phagocytosis, and inflammasome activation, which are the most well-characterized ways that autophagy proteins regulate inflammation. In addition to the increase in proinflammatory cytokine production during *M. tuberculosis* infection, loss of ATG5 in innate immune cells also results in an early induction of T_H_17 responses. Despite prior published *in vitro* cell culture experiments supporting a role for autophagy in controlling *M. tuberculosis* replication in macrophages, loss of autophagy does not affect *M. tuberculosis* burden in macrophages *in vivo* and, therefore, the effects of autophagy on inflammatory responses occur without changes in pathogen numbers. These findings reveal new roles for autophagy proteins in lung resident macrophages and dendritic cells that are required to suppress inflammatory responses that are associated with poor control of *M. tuberculosis* infection.

## INTRODUCTION

According to the World Health Organization, 10 million people fell ill with *Mycobacterium tuberculosis* infection and 1.5 million people died of tuberculosis (TB) in 2020, marking the first increase in TB-associated deaths in over a decade (1). Whether a person controls the initial *M. tuberculosis* infection or develops active TB disease is directly impacted by the type of immune response elicited in the infected individual (2). Therefore, better understanding of what constitutes a protective versus non-protective immune response to *M. tuberculosis* infection is critical for developing better therapies and prevention measures to fight this deadly disease. Genetic mouse models have provided invaluable insight into the immunological processes that are required for control of *M. tuberculosis* infection. Infection of mice through the aerosol route leads to phagocytosis of *M. tuberculosis* by alveolar macrophages, initiating an inflammatory response and recruitment of innate immune cells to the lung (2). *M. tuberculosis* replicates within these innate immune cells until antigen specific T cells traffic to the lung where they activate the innate immune cells to restrain *M. tuberculosis* replication and suppress inflammation. *M. tuberculosis* establishes a chronic infection in wild-type (WT) mice, which survive for over a year with this infection.

*Atg5^fl/fl^-LysM-Cre* mice, which delete the *Atg5* gene specifically in macrophages, inflammatory monocytes, some dendritic cells (DCs), and neutrophils, are severely susceptible to *M. tuberculosis* infection (3–5), highlighting ATG5 as a critical component of a protective immune response to *M. tuberculosis*. *M. tuberculosis* infected *Atg5^fl/fl^-LysM-Cre* mice fail to control bacterial replication and succumb to infection by 40 days post-infection (dpi) (3–5). The uncontrolled *M. tuberculosis* replication is associated with an early (by 14 dpi) and sustained influx of neutrophils in the *M. tuberculosis* infected *Atg5^fl/fl^-LysM-Cre* mice. Depletion of neutrophils during *M. tuberculosis* infection in *Atg5^fl/fl^-LysM-Cre* mice rescues the susceptibility and extends their survival (3), demonstrating that the neutrophil-dominated inflammation contributed to their susceptibility. In general, higher abundance of neutrophils during *M. tuberculosis* infection have been associated with worse disease outcomes in mice (6–13) and humans (12,14–17). Therefore, understanding the regulatory mechanisms that govern neutrophil recruitment and accumulation during *M. tuberculosis* infection could be key for manipulating inflammatory responses to better control TB.

ATG5 is required for the intracellular pathway of autophagy, a process by which cytoplasmic contents are targeted to the lysosome for degradation (18,19). Initiation of autophagy involves phagophore formation from the endoplasmic reticulum, which is mediated by the ULK1 complex (ULK1/ULK2, ATG13, FIP200, ATG101) and the PI3 kinase complex (ATG14L, BECLIN1, VPS15, and VPS34)(20,21). Elongation of the autophagosomal double membrane depends on two ubiquitin-like conjugation systems. In the first system, ATG12 is activated by ATG7, transferred to ATG10, and covalently attached to ATG5. The second ubiquitin-like component is LC3 (microtubule-associated protein 1 light chain 3), which is conjugated to phosphatidylethanolamine, generating the membrane bound form called LC3-II through the actions of ATG7 and ATG3. ATG5-ATG12 facilitates LC3 lipidation through its interactions with ATG3, while ATG16L1 specifies the localization of LC3 conjugation to the autophagosome membrane (18,19,21,22). The autophagosome membrane is then completed and targeted for fusion with the lysosome where the autophagosome cargo are degraded. In addition, ATG5 also functions outside of autophagy, including during *M. tuberculosis* infection (3), although these activities remain poorly understood. Recent work supports an autophagy-dependent role for ATG5 in LysM^+^ innate immune cells in suppressing neutrophil recruitment to the lungs during *M. tuberculosis* infection (23). Using *in vitro* cell culture experiments, multiple groups have reported that macrophages require autophagy to control *M. tuberculosis* replication by targeting the pathogen to the lysosome (xenophagy) as well as to prevent necrosis following days of infection in culture (5,23–29). However, to date there is no evidence that xenophagy functions in this capacity *in vivo*. Therefore, the mechanistic basis for how loss of ATG5 results in early and exagerated recruitment of neutrophils during *M. tuberculosis* infection *in vivo* remains unknown.

In this manuscript, we dissect the role for ATG5 in regulating neutrophil recruitment and accumulation during *M. tuberculosis* infection *in vivo*. We find that ATG5 functions with other autophagy proteins specifically in CD11c^+^ lung macrophages and DCs to limit the production of cytokines and chemokines that otherwise promote neutrophil influx to the lung early in *M. tuberculosis* infection. We demonstrate that loss of autophagy in macrophages and DCs does not affect *M. tuberculosis* burden in these cell types *in vivo* and instead changes the inflammatory response to the infection at time points before cell survival is compromised. In addition, ATG5 is required in lung macrophages and DCs to limit IL-17A production from CD4^+^ T cells. Together, our studies reveal new roles for ATG5 and other autophagy proteins in regulating inflammatory responses during infection, which with further dissection could provide insight into pathways that may be targeted to effectively promote protective immune responses during TB.

## RESULTS

### ATG5 is required in CD11c^+^ lung macrophages and DCs to control neutrophil recruitment and accumulation early during *M. tuberculosis* infection *in vivo*

*M. tuberculosis* infection of *Atg5^fl/fl^-LysM-Cre* mice results in the recruitment of a higher number of neutrophils in the lungs at 14 dpi as compared to *Atg5^fl/fl^*controls, despite equivalent bacterial burdens at this time point (3). There are also no differences in the abundance of non-neutrophil cell types in *Atg5^fl/fl^-LysM-Cre* and *Atg5^fl/fl^* mice at 14 dpi (3). This indicates that specifically neutrophils are accumulating in *M. tuberculosis*-infected *Atg5^fl/fl^-LysM-Cre* mice due to a defect in inflammatory responses to infection and not due to higher burden. To determine which LysM^+^ cells required ATG5 to control the early influx of neutrophils into the lungs during *M. tuberculosis* infection, we compared bacterial burdens and neutrophil inflammation in *Atg5^fl/fl^-LysM-Cre*, *Atg5^fl/fl^-Mrp8-Cre* (deletion in neutrophils)*, Atg5^fl/fl^-Cd11c-Cre* (deletion in lung macrophages and DCs), and *Atg5^fl/fl^* controls at 14 dpi. At 14 dpi, the *Atg5^fl/fl^-LysM-Cre* and *Atg5^fl/fl^-Cd11c-Cre* mice, but not *Atg5^fl/fl^-Mrp8-Cre* mice, had higher levels of neutrophil inflammation in the lungs as compared to *Atg5^fl/fl^* controls **(Fig 1A and 1B)**. The degree of increased neutrophil frequency was similar in *Atg5^fl/fl^-LysM-Cre* and *Atg5^fl/fl^-Cd11c-Cre* mice, indicating that loss of *Atg5* in CD11c^+^ cells, but not neutrophils, leads to the early influx of neutrophils into the lungs during *M. tuberculosis* infection. At 14 dpi, none of the mouse strains harbored increased *M. tuberculosis* burden in their lungs **(Fig 1C)**, indicating that the increase in neutrophil abundance in *Atg5^fl/fl^-Cd11c-Cre* mice is not due to elevated bacterial burden and reflects a dysregulated inflammatory response to infection.

**Figure 1.**
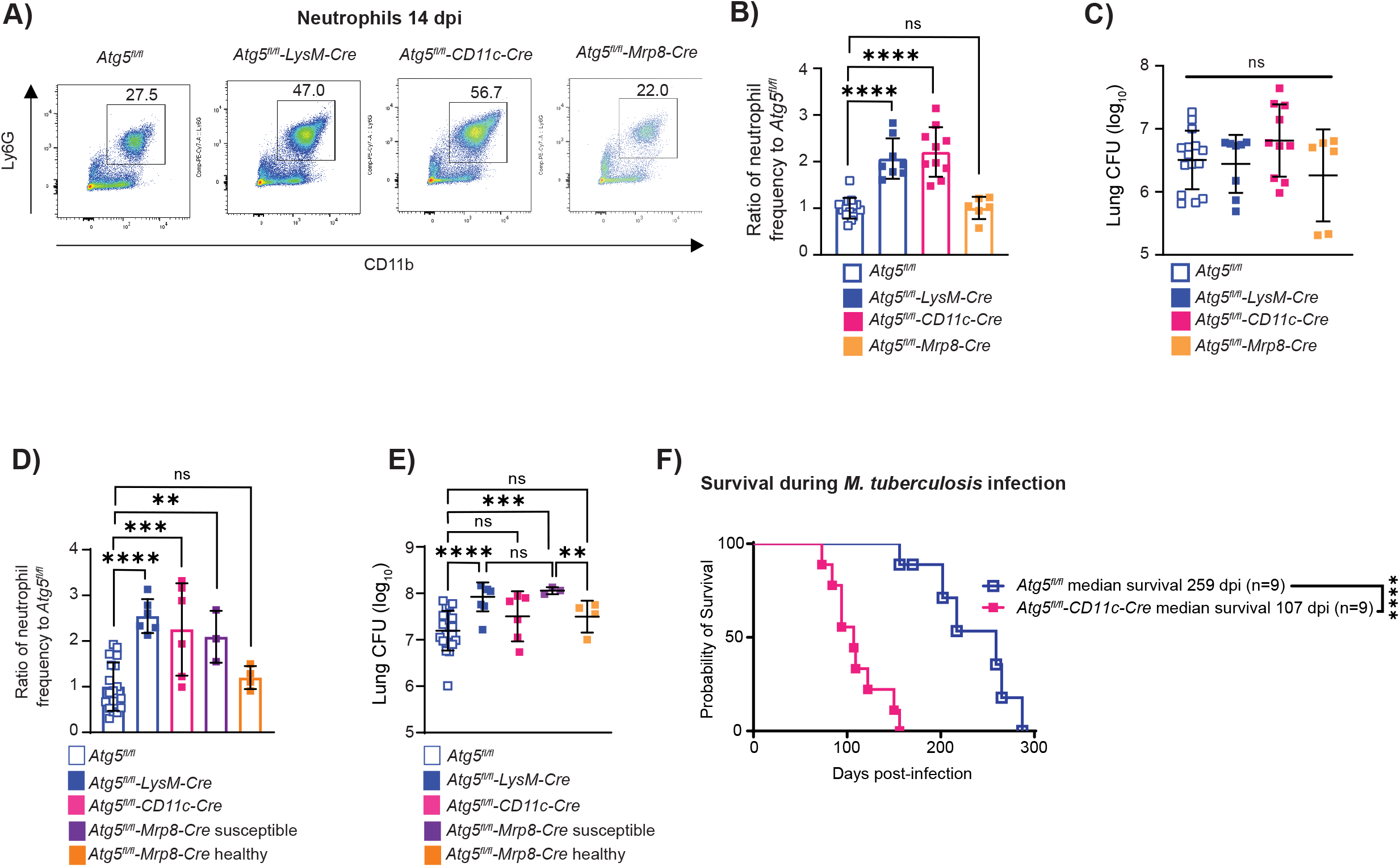
ATG5 is required in CD11c^+^ cells to regulate the early influx of neutrophils during *M. tuberculosis* infection *in vivo*. **(A)** Representative flow cytometry plots of neutrophils (CD45^+^Ly6G^+^CD11b^+^) at 14 days post-infection (dpi) from *Atg5^fl/fl^, Atg5^fl/fl^-LysM-Cre, Atg5^fl/fl^-CD11c-Cre* and *Atg5^fl/fl^-Mrp8-Cre* mice. **(B)** Proportion of CD45^+^ cells that are neutrophils in the lung at 14 dpi in *Mtb-*GFP infected *Atg5^fl/fl^* (n=15), *Atg5^fl/fl^-LysM-Cre* (n=8)*, Atg5^fl/fl^-CD11c-Cre* (n=10), and *Atg5^fl/fl^-Mrp8-Cre* (n=6) mice. Neutrophil frequency is reported as a ratio relative to the average neutrophil frequency in *Atg5^fl/fl^* control mice at 14 dpi within a given experiment. **(C)** Lung burden from the right lung at 14 dpi in *Mtb-*GFP infected *Atg5^fl/fl^* (n=15)*, Atg5^fl/fl^-LysM-Cre* (n=8)*, Atg5^fl/fl^-CD11c-Cre* (n=10), and *Atg5^fl/fl^-Mrp8-Cre* (n=6) mice. **(D)** Proportion of CD45^+^ cells that are neutrophils in the lung at 21 dpi in *Mtb-*GFP infected *Atg5^fl/fl^*(n=19)*, Atg5^fl/fl^-LysM-Cre* (n=6)*, Atg5^fl/fl^-CD11c-Cre* (n=6) and *Atg5^fl/fl^-Mrp8-Cre* (n=7) mice. Neutrophil frequency is reported as a ratio relative to the average neutrophil frequency in *Atg5^fl/fl^* control mice at 21 dpi within a given experiment. Susceptible and healthy *Atg5^fl/fl^-Mrp8-Cre* mice are defined as done previously where susceptible *Atg5^fl/fl^-Mrp8-Cre* mice have lost more than 5% of their pre-infection body weight by 20 dpi and healthy *Atg5^fl/fl^-Mrp8-Cre* mice have lost less than 5% of their pre-infection body weight at 20 dpi (3). **(E)** Lung burden from the right lung at 21 dpi in *Mtb-*GFP infected *Atg5^fl/fl^* (n=19)*, Atg5^fl/fl^-LysM-Cre* (n=6)*, Atg5^fl/fl^-CD11c-Cre* (n=6), and *Atg5^fl/fl^-Mrp8-Cre* (n=7) mice. **(F)** Kaplan Meier curve of survival proportions during *Mtb-*GFP infection of *Atg5^fl/fl^* (n=9) and *Atg5^fl/fl^-CD11c-Cre* (n=9) mice. Statistical differences were determined by a log-rank Mantel-Cox test (F) or one-way ANOVA and Šídák multiple comparison test (B-E). * P < 0.05, ** P < 0.01, *** P < 0.001, **** P < 0.0001. Differences that are not statistically significant are designated as ns. Pooled data from at least two separate experiments is graphed where each data point is from one biological replicate.

To determine if the higher levels of neutrophils in the lungs of *M. tuberculosis*-infected *Atg5^fl/fl^-CD11c-Cre* mice was due to elevated neutrophil abundance in circulation prior to or during infection, we monitored neutrophil frequency in the blood in uninfected and 14 dpi *Atg5^fl/fl^* and *Atg5^fl/fl^-CD11c-Cre* mice. There was no significant difference in the frequency of neutrophils in the blood of uninfected or 14 dpi *Atg5^fl/fl^* and *Atg5^fl/fl^-CD11c-Cre* mice **(S1A Fig)**, suggesting that the accumulation of neutrophils in the lungs of *M. tuberculosis*-infected *Atg5^fl/fl^-CD11c-Cre* mice was due to specific recruitment of neutrophils to the site of infection or an inability to clear neutrophils from the lung. To begin to investigate this latter possibility, we monitored whether dead neutrophils were accumulating in the lungs of *Atg5^fl/fl^-CD11c-Cre* mice during *M. tuberculosis* infection by analyzing neutrophil viability at 14 dpi by flow cytometry **(S1B Fig)**. We did not observe a significant difference in the frequency of viable neutrophils at 14 dpi in *Atg5^fl/fl^* and *Atg5^fl/fl^-CD11c-Cre* mice, indicating that increased neutrophil inflammation in the lungs of *Atg5^fl/fl^-CD11c-Cre* mice was not due to differences in neutrophil viability.

The higher levels of neutrophils in the lungs of *Atg5^fl/fl^-LysM-Cre* and *Atg5^fl/fl^-Cd11c-Cre* mice were sustained through 21 dpi **(Fig 1D)**. However, only *Atg5^fl/fl^-LysM-Cre* mice, and not *Atg5^fl/fl^-Cd11c-Cre* mice, had higher bacterial burdens in the lungs at 21 dpi **(Fig 1E)**, similar to as previously reported (3). Loss of *Atg5* in neutrophils results in increased susceptibility to *M. tuberculosis* infection in some, but not all, *Atg5^fl/fl^-Mrp8-Cre* mice (3). The susceptible *Atg5^fl/fl^-Mrp8-Cre* mice accumulate higher neutrophil numbers and bacterial burdens in their lungs at 21 dpi **(Fig 1D and 1E)** (3). Therefore, loss of *Atg5* in neutrophils is likely contributing to the higher burdens in the lungs of *Atg5^fl/fl^-LysM-Cre* mice at 21 dpi. These data indicate that ATG5 has a role in CD11c^+^ lung macrophages and DCs to regulate early recruitment of neutrophils, but not the control of *M. tuberculosis* replication during *M. tuberculosis* infection at 14 and 21 dpi.

To determine how the loss of Atg5 in CD11c^+^ cells and the resulting early influx of neutrophils into the lungs affected host susceptibility, we monitored survival in *M. tuberculosis* infected *Atg5^fl/fl^-Cd11c-Cre* mice as compared to *Atg5^fl/fl^* controls. *Atg5^fl/fl^-Cd11c-Cre* mice succumbed to *M. tuberculosis* infection between 100 and 150 dpi, which was significantly earlier than *Atg5^fl/fl^* controls (median survival time of 259 dpi) **(Fig 1F)**, but not as early as *Atg5^fl/fl^-LysM-Cre* mice (succumb 30-40 dpi (3)). These data demonstrate that ATG5 is required in CD11c^+^ lung macrophages and DCs to control early neutrophil recruitment and promote survival following *M. tuberculosis* infection.

### The role for ATG5 in lung macrophages and DCs in regulating neutrophil recruitment is dependent on other autophagy proteins

Deletion of multiple different autophagy genes in all LysM^+^ innate immune cells can result in increased neutrophil recruitment to the lung during *M. tuberculosis* infection (23). However, we previously showed that at least one role for ATG5 in LysM^+^ innate immune cells in controlling *M. tuberculosis* infection is autophagy-independent (3). Therefore, it is not known if the role for ATG5 specifically in CD11c^+^ lung macrophages and DCs is the same as described when broadly deleting *Atg5* in all LysM^+^ cells. To determine whether the regulation of neutrophil recruitment by ATG5 in CD11c^+^ lung macrophages and DCs was dependent on other autophagy proteins or represented the autophagy-independent role for ATG5 during *M. tuberculosis* infection, we monitored neutrophil abundance in the lungs of mice lacking expression of another essential autophagy protein, BECLIN 1, in CD11c^+^ cells (*Becn1^fl/fl^-Cd11c-Cre*) at 14 dpi by flow cytometry. Similar to *Atg5^fl/fl^-Cd11c-Cre* mice, *Becn1^fl/fl^-Cd11c-Cre* mice also exhibited elevated neutrophil frequency in the lung at 14 dpi relative to *Becn1^fl/fl^* control mice **(Fig 2A)**, despite no difference in bacterial burden **(Fig 2B)**. In addition, analysis of *M. tuberculosis* infected *Atg16l1^fl/fl^-LysM-Cre* and *Becn1^fl/fl^-LysM-Cre* mice also revealed higher levels of neutrophils in the lungs at 14 dpi relative to controls, without higher bacterial burdens **(Fig 2C and 2D)**.

**Figure 2.**
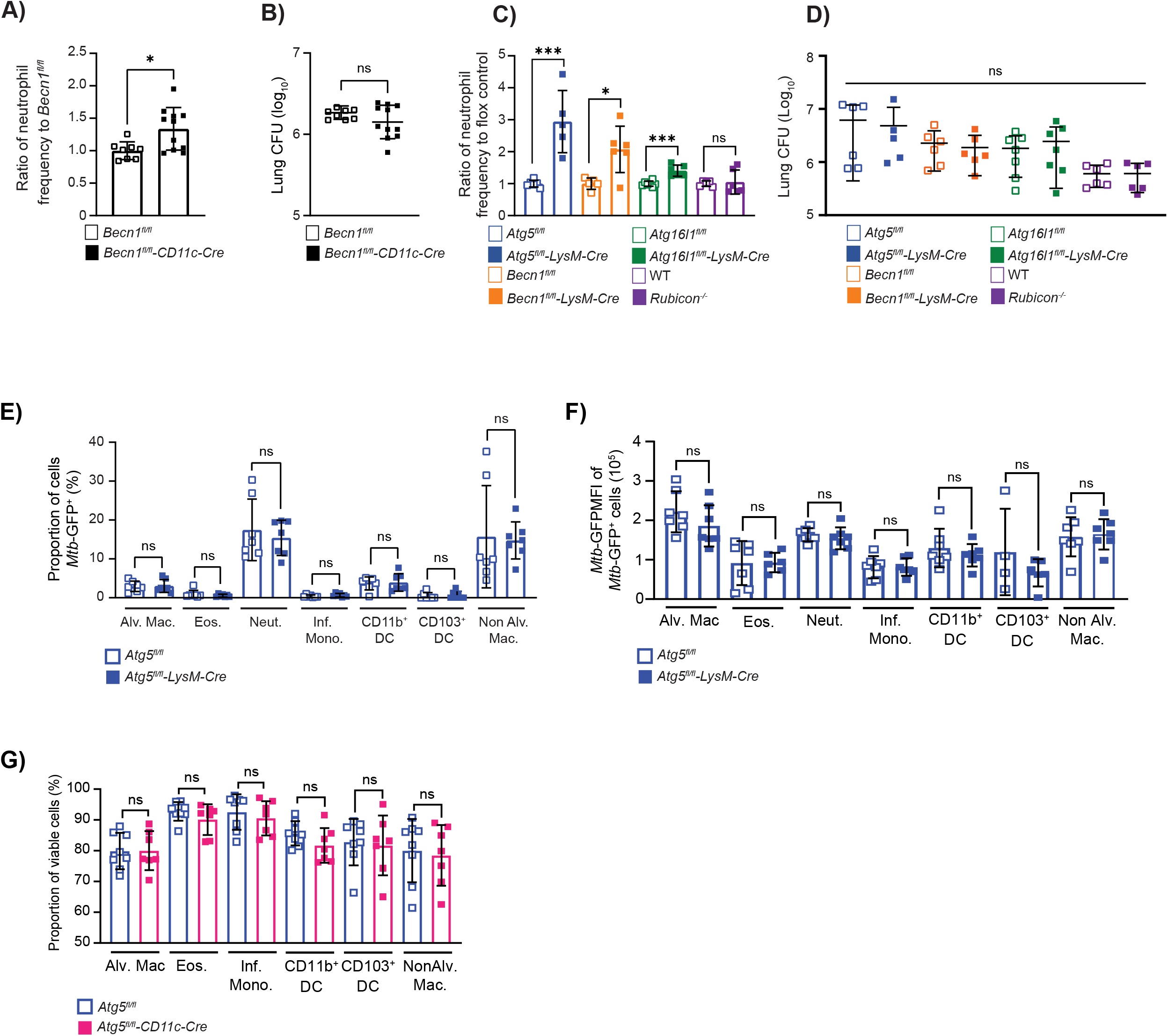
The role for ATG5 in lung macrophages and DCs in regulating neutrophil recruitment is dependent on other autophagy proteins but does not involve control of pathogen infection or burden. **(A)** Proportion of CD45^+^ cells that are neutrophils (CD45^+^Ly6G^+^CD11b^+^) in the lung at 14 dpi in *Mtb-*GFP infected *Becn1^fl/fl^* (n=8) or *Becn1^fl/fl^-CD11c-Cre* (n=11) mice. Neutrophil frequency is reported as a ratio relative to the average neutrophil frequency in *Becn1^fl/fl^*control mice at 14 dpi. **(B)** Lung burden from the right lobes of the lung at 14 dpi in *Mtb-*GFP infected *Becn1^fl/fl^* (n=8) or *Becn1^fl/fl^-CD11c-Cre* (n=11) mice. **(C)** Neutrophil frequency of CD45^+^ cells reported as a ratio to the average neutrophil frequency in floxed control mice at 14 dpi in *Mtb-*GFP infected *Atg5^fl/fl^* (n=6), *Atg5^fl/fl^-LysM-Cre* (n=5), *Becn1^fl/fl^* (n=6), *Becn1^fl/fl^-LysM-Cre* (n=6), *Atg16l1^fl/fl^* (n=7), *Atg16l1^fl/fl^-LysM-Cre* (n=7) mice. Neutrophil frequencies at 14 dpi in *Mtb-*GFP infected *Rubicon^-/-^* (n=6) mice were compared to wildtype C57BL/6J (n=6) mice. **(D)** Lung burden from right lobes of the lung at 14 dpi in *Mtb-*GFP infected *Atg5^fl/fl^*(n=6), *Atg5^fl/fl^-LysM-Cre* (n=5), *Becn1^fl/fl^*(n=6), *Becn1^fl/fl^-LysM-Cre* (n=6), *Atg16l1^fl/fl^*(n=7), *Atg16l1^fl/fl^-LysM-Cre* (n=7), wildtype C57BL/6J (n=6) and *Rubicon^-/-^* (n=6) mice. **(E,F)** The proportion of *M. tuberculosis* infected (*Mtb-*GFP*^+^*) and the median fluorescence intensity of *Mtb-*GFP in infected alveolar macrophage (Alv. Mac.), eosinophils (Eos.), neutrophils (Neut.), inflammatory monocytes (Inf. Mono.), CD11b^+^ DC, CD103^+^ DC, and non-alveolar macrophages in the lung in *Atg5^fl/fl^* (n=7) and *Atg5^fl/fl^-LysM-Cre* (n=7) mice at 14 dpi. **(G)** The proportion of viable (Zombie^-^) alveolar macrophages, eosinophils, neutrophils, inflammatory monocytes, CD11b^+^ DC, CD103^+^ DC and non-alveolar macrophages in the lung in *Atg5^fl/fl^* (n=8) and *Atg5^fl/fl^-CD11c-Cre* (n=7) mice at 14 dpi. Statistical differences were determined by student t-test to compare the LysM-Cre or CD11c-Cre mice to their respective floxed control and *Rubicon^-/-^*mice to wildtype C57BL/6J mice (A-G).* P < 0.05, ** P < 0.01, *** P < 0.001, **** P < 0.0001. Differences that are not statistically significant are designated as ns. Pooled data from at least two separate experiments is graphed where each data point is from one biological replicate.

In addition to their role in canonical autophagy, the proteins ATG5, BECLIN 1, and ATG16L1 are also required for the process of LC3 associated phagocytosis (LAP), where LC3 is recruited to the phagosome, resulting in LC3^+^ single membrane vesicles that traffic to the lysosome for degradation. LAP can dampen inflammatory responses through efferocytosis, pathogen removal, stimulating inhibitory immune-receptor signaling, and reducing auto-antigen levels (30–33). In contrast to canonical autophagy, LAP uses RUBICON and UVRAG instead of ATG14 in the PI3K complex and does not depend on ULK1 (33,34). To distinguish between whether ATG5, BECLIN 1 and ATG16L1 were functioning through autophagy or LAP to regulate neutrophil recruitment during *M. tuberculosis* infection, we infected mice lacking RUBICON expression, a protein specifically required for LAP. *Rubicon*^-/-^ mice had no difference in neutrophil accumulation or bacterial burdens as compared to WT controls following *M. tuberculosis* infection **(Fig 2C and 2D)**, indicating that LAP is not required to control neutrophil inflammation during *M. tuberculosis* infection. Importantly, BECLIN 1 and ATG5 function at different steps of autophagy. Therefore, the requirement of both BECLIN 1 and ATG5 suggests that both the initiation and elongation steps of autophagy are required in CD11c^+^ cells to control neutrophil recruitment early during *M. tuberculosis* infection. Targeting of pathogens to the lysosome via autophagy is termed xenophagy and multiple studies have reported roles for xenophagy in controlling *M. tuberculosis* replication in macrophages and DCs in cell culture *in vitro* (5,23–29). However, in addition to there being no differences in bacterial burden in the lungs at 14 dpi **(Fig 1C and 2D)** (3), there was no significant difference in the proportion of macrophages, eosinophils, neutrophils, inflammatory monocytes, or DCs that were infected with *M. tuberculosis* **(Fig 2E)** and no difference in the *M. tuberculosis* burden in autophagy-deficient cells in the lungs of *Atg5^fl/fl^-LysM-Cre* and *Atg5^fl/fl^* mice at 14 dpi **(Fig 2F).** Loss of xenophagy in bone marrow derived macrophages *in vitro* has also been associated with increased necrosis during *M. tuberculosis* infection (23). However, we did not detect a difference in the viability of macrophages, inflammatory monocytes, eosinophils, neutrophils or DCs in the lungs of *Atg5^fl/fl^* and *Atg5^fl/fl^-Cd11c-Cre* mice at 14 dpi **(Fig 2G, S1B Fig)**, although we cannot rule out effects on the balance of different cell death pathways. These data support that the role for autophagy in CD11c^+^ lung macrophages and DCs during *M. tuberculosis* infection *in vivo* is independent of xenophagy regulating *M. tuberculosis* replication. In addition, these data show that the elevated neutrophil abundance in *M. tuberculosis*-infected *Atg5^fl/fl^-LysM-Cre* mice is not due to differences in the cell viability or the cell types infected with *M. tuberculosis* but instead is driven by an imbalanced inflammatory response.

### Autophagy regulates proinflammatory responses in macrophages during *M. tuberculosis* infection

We have previously shown that lungs of *Atg5^fl/fl^-LysM-Cre* mice at 14 dpi contain higher levels of G-CSF and IL-17A than control mice (3), cytokines that promote neutrophil development and recruitment. At this time point, the primary CD11c^+^ cell types that are infected by *M. tuberculosis* are the lung macrophages, encompassing alveolar and interstitial macrophages (35,36). Therefore, we hypothesized that autophagy could be suppressing the production of these cytokines from *M. tuberculosis* infected macrophages. We tested this hypothesis by culturing bone marrow derived macrophages (BMDMs) from *Atg5^fl/fl^, Atg5^fl/fl^-LysM-Cre, Atg16l1^fl/fl^, Atg16l1^fl/fl^-LysM-Cre, Becn1^fl/fl^* and *Becn1^fl/fl^-LysM-Cre* mice and infecting with *M. tuberculosis in vitro* before monitoring cytokine and chemokine production using a cytokine bead array (Bio-Rad) on the supernatants from infected cultures **(Fig 3A-3F and S2 Fig)**. Of the 23 cytokines tested, we detected significantly higher levels of IL-1β, G-CSF, KC, TNF-α and RANTES from the *Atg5^fl/fl^-LysM-Cre* macrophage cultures compared to controls at 24 hpi **(Fig 3A-3E)**, despite no difference in bacterial burdens or BMDM viability at this time point **(Fig 3F and S2A Fig)**, indicating that the heightened proinflammatory response of *Atg5^-/-^*BMDMs was not in response to increased antigen or macrophage cell death. The levels of these cytokines were only different following *M. tuberculosis* infection and not in mock infected cultures, indicating that the increased pro-inflammatory responses were infection-induced. The higher levels of G-CSF and KC, both pro-inflammatory signals associated with neutrophil inflammation (37–39), produced from *Atg5*-deficient macrophages in response to *M. tuberculosis* infection were dose dependent and confirmed by ELISA **(Fig 3H and 3I)**. Similar to *Atg5^fl/fl^-LysM-Cre* BMDMs, *Atg16l1^fl/fl^-LysM-Cre* BMDMs also produced higher levels of IL-1β, G-CSF and TNF-α following *M. tuberculosis* infection *in vitro* **(Fig 3A, 3B and 3D)**. *Becn1^fl/fl^-LysM-Cre* BMDMs also produced more IL-1β and G-CSF following *M. tuberculosis* infection *in vitro* compared to control cells (**Fig 3A, 3B and S2 Fig**) despite no difference in *M. tuberculosis* burden at this time point (**Fig 3F**). In addition, *Becn1^fl/fl^-LysM-Cre* BMDMs produced higher levels of IL-6, MIP-1α, MIP-1β, and MCP-1 following *M. tuberculosis* infection (**Fig 3G and S2 Fig**). IL-6 in particular is associated with neutrophil recruitment (40–43) and also trended higher in *M. tuberculosis*-infected *Atg5^fl/fl^-LysM-Cre* BMDMs **(Fig 3G)**, so we further analyzed the levels of IL-6 produced by *M. tuberculosis*-infected *Atg5^fl/fl^-LysM-Cre* BMDMs using an ELISA **(Fig 3J)**. These analyses revealed a dose-dependent increase of IL-6 secretion in *M. tuberculosis*-infected *Atg5^fl/fl^-LysM-Cre* BMDMs. We were able to detect more cytokines and chemokines being differentially produced by *M. tuberculosis* infected BMDMs than what was previously detected in the lungs of *Atg5^fl/fl^-LysM-Cre* mice at 14dpi (3). This is likely due to the dilution of signals produced by macrophages in the context of the total lung homogenate, decreasing our sensitivity to detect macrophage specific responses. In addition, it is possible that the inflammatory responses of BMDMs differ from lung macrophages and DCs during *M. tuberculosis* infection. Nonetheless, loss of expression of the autophagy proteins ATG5, ATG16L1, or BECLIN 1 in macrophages both *in vivo* and *in vitro* results in higher levels of cytokines and chemokines that are associated with neutrophil recruitment and accumulation following *M. tuberculosis* infection relative to controls, indicating that canonical autophagy is required in macrophages to control proinflammatory responses during *M. tuberculosis* infection without affecting *M. tuberculosis* burden in macrophages.

**Figure 3.**
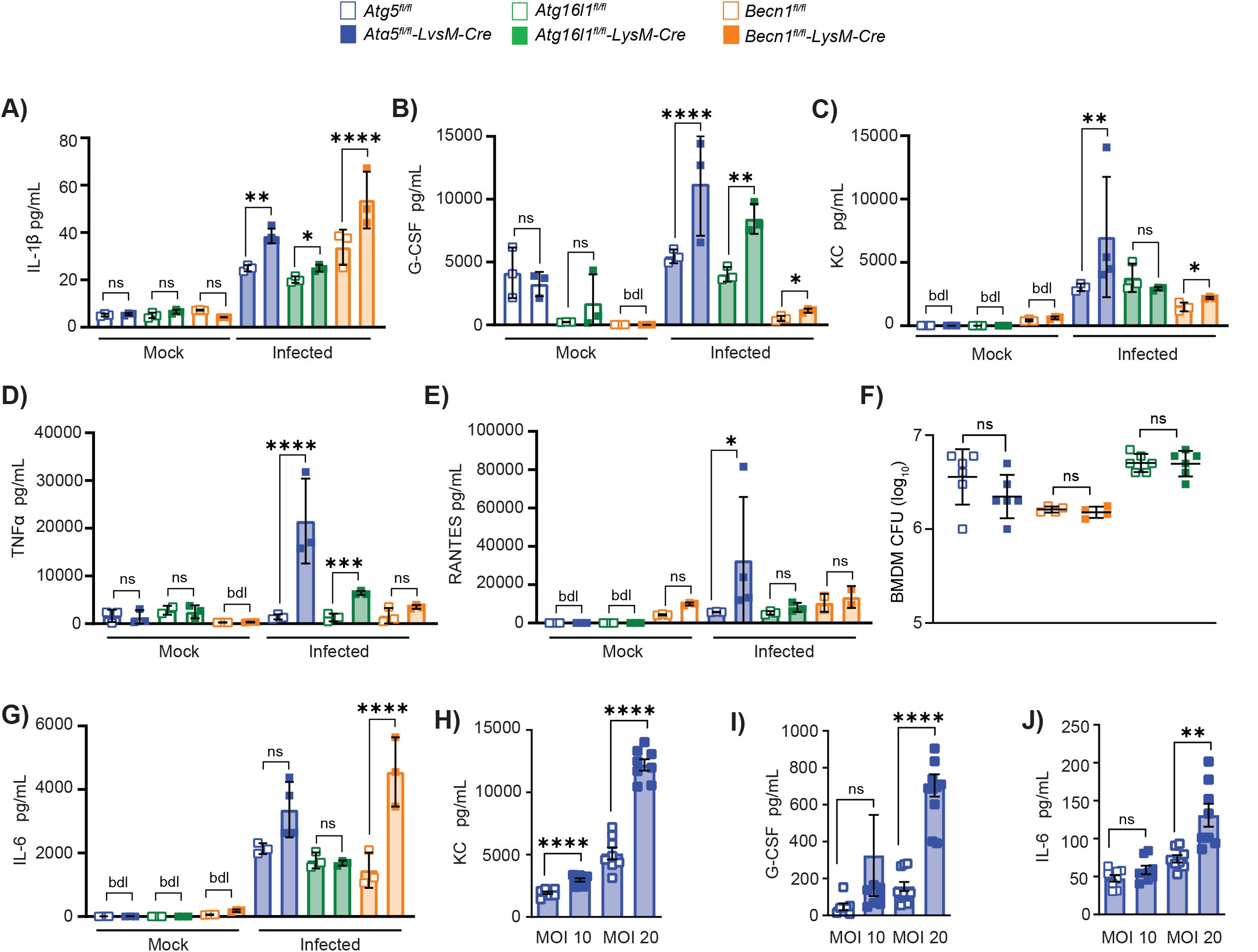
Autophagy regulates proinflammatory responses in macrophages during *M. tuberculosis* infection. **(A)** Cytokine bead array data to quantify cytokines in culture supernatants from *Atg5^fl/fl^, Atg5^fl/fl^-LysM-Cre^-^, Atg16l1^fl/fl^, Atg16l1^fl/fl^-LysM-Cre, Becn1^fl/fl^*or *Becn1^fl/fl^-LysM-Cre* BMDMs mock-treated or infected with *Mtb-*GFP at an MOI of 10 for 24 hours. BMDMs generated from at least 3 mice were tested in duplicate to quantify cytokine production. **(A)** IL-1β, **(B)** G-CSF, **(C)** KC, **(D)** TNF-α, and **(E)** RANTES levels at 24 hpi are shown. **(F)** BMDM CFU counts at 24 hpi. **(G)** IL-6 levels at 24 hpi are shown. **(H)** KC, **(I)** G-CSF, and **(J)** IL-6 levels at 24 hpi in *Atg5^fl/fl^* (n=8) and *Atg5^fl/fl^-LysM-Cre* (n=8) BMDMs infected with *M. tuberculosis* at an MOI of 10 or 20 for 24 hours determined by ELISA. Statistical differences were determined by student t-test to compare the autophagy gene-deficient cells to their respective floxed control cells (A-J). * P < 0.05, ** P < 0.01, *** P < 0.001, **** P < 0.0001. Differences that are not statistically significant are designated as ns. Cytokine levels below detection limits are designated as dbl. Each data point is one biological replicate, and the samples were generated from at least two separate experiments.

### Autophagy suppresses neutrophil recruitment early during *M. tuberculosis* infection independent of mitophagy and inflammasome activation

Autophagy has been shown to suppress proinflammatory responses by negatively regulating inflammasome activation indirectly through regulation of NFκB signaling and directly by degrading pro-IL-1β and clearance of inflammasome components (44–46), which can otherwise promote pro-inflammatory responses, IL-1β secretion and neutrophil recruitment (44,45,47–49). Indeed, *Atg5^fl/fl^-LysM-Cre*, *Atg16l1^fl/fl^-LysM-Cre* and *Becn1^fl/fl^-LysM-Cre* BMDMs produce significantly more IL-1β in response to *M. tuberculosis* infection *in vitro* at 24 hpi compared to control BMDMs (**Fig 3A**), supporting that loss of autophagy has resulted in increased inflammasome activation. The primary inflammasome activated during *M. tuberculosis* infection of macrophages is the NLRP3 inflammasome, which consists of the NOD-, LRR- and pyrin-domain containing protein 3 (NLRP3) sensor, ASC adaptor, and CASPASE 1(50–53). TLR engagement and NFκB activation during *M. tuberculosis* infection constitute the priming step of inflammasome activation, resulting in increased expression of pro-IL-1β and NLRP3 (54,55). Phagocytosis of *M. tuberculosis* and subsequent Esx-1-dependent plasma membrane damage leading to potassium efflux is the second signal promoting NLRP3 inflammasome formation, which mediates CASPASE 1 activation followed by IL-1β processing and secretion (26, 29).

To determine whether the increased neutrophil inflammation following *M. tuberculosis* infection in autophagy-deficient mice results from increased inflammasome activation, we crossed *Caspase1/11^-/-^*mice to *Atg5^fl/fl^-LysM-Cre* and *Becn1^fl/fl^-LysM-Cre* mice and monitored neutrophil abundance in the lungs at 14 dpi. *Caspase1/11^-/-^/Atg5^fl/fl^-LysM-Cre and Caspase1/11^-/-^/Becn1^fl/fl^-LysM-Cre* mice had similar neutrophil abundances and bacterial burdens in the lungs at 14 dpi as *Atg5^fl/fl^-LysM-Cre mice* and *Becn1^fl/fl^-LysM-Cre* mice, respectively **(Fig 4A and 4B)**, indicating that increased neutrophil recruitment in the absence of autophagy is occurring independent of CASPASE1/11. *Caspase1/11* deletion also did not extend the survival of *Atg5^fl/fl^-LysM-Cre* mice, indicating that increased inflammasome activation does not contribute to the early susceptibility of these mice **(Fig 4C)**.

**Figure 4.**
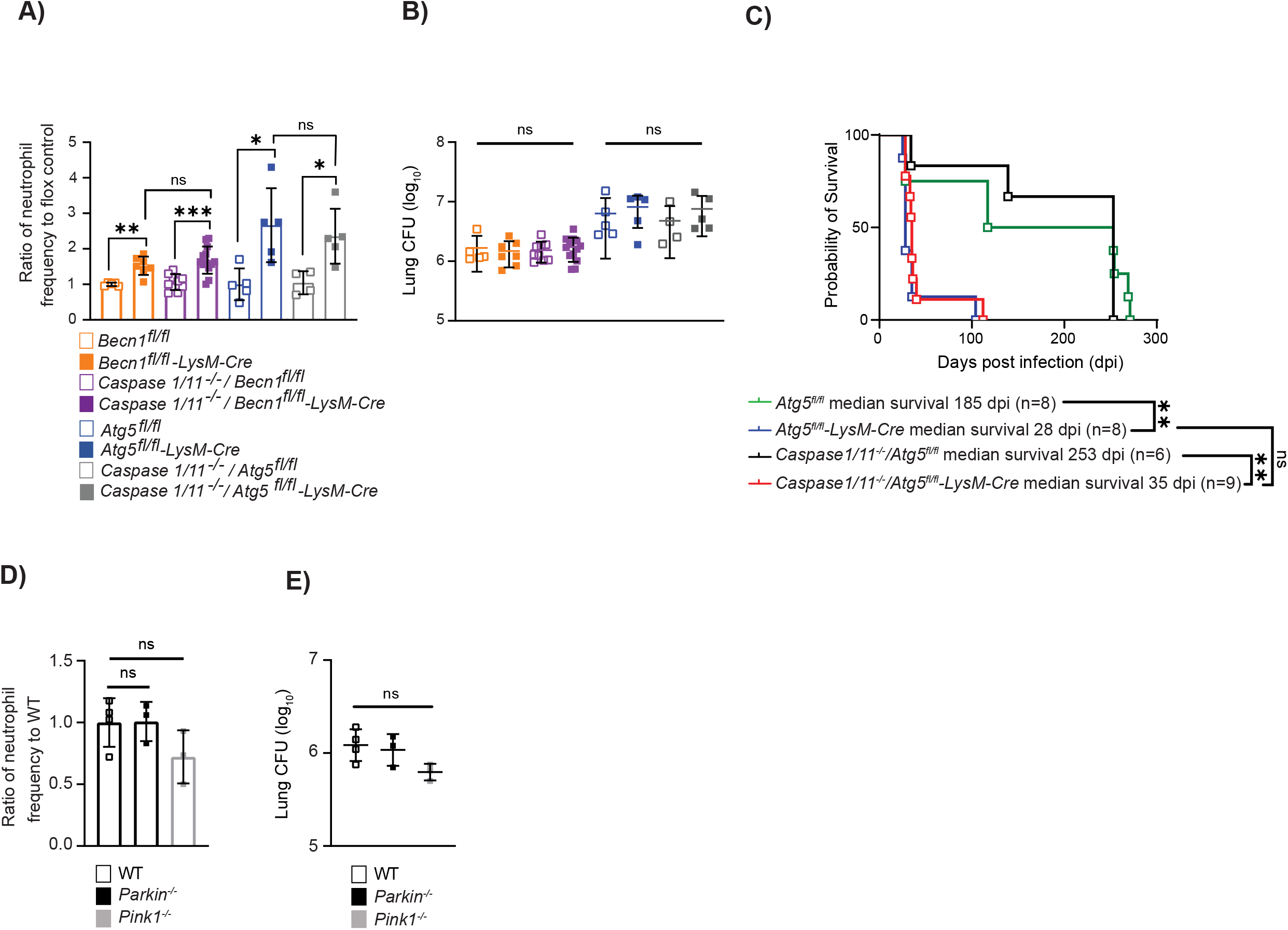
Autophagy suppresses neutrophil recruitment independent of mitophagy and inflammasome activation during *M. tuberculosis* infection. **(A)** Proportion of CD45^+^ cells that are neutrophils (CD45^+^Ly6G^+^CD11b^+^) in the lung at 14 dpi in *Mtb-*GFP infected *Becn1^fl/fl^* (n=5), *Becn1^fl/fl^-LysM-Cre* (n=7), *Caspase1/11^-/-^/Becn1^fl/fl^* (n=9), *Caspase1/11^-/-^/Becn1^fl/fl^-LysM-Cre* (n=13), *Atg5^fl/fl^* (n=5), *Atg5^fl/fl^-LysM-Cre* (n=5), *Caspase1/11^-/-^/Atg5^fl/fl^*(n=4), or *Caspase1/11^-/-^/Atg5^fl/fl^-LysM-Cre* (n=5) mice reported as a ratio relative to the average neutrophil frequency in corresponding floxed control mice. **(B)** Lung burden at 14 dpi from right lobes of the lung in *Mtb-*GFP infected *Becn1^fl/fl^* (n=5), *Becn1^fl/fl^-LysM-Cre* (n=7), *Caspase1/11^-/-^/Becn1^fl/fl^*(n=9), *Caspase1/11^-/-^/Becn1^fl/fl^-LysM-Cre* (n=13), *Atg5^fl/fl^* (n=5), *Atg5^fl/fl^-LysM-Cre* (n=5), *Caspase1/11^-/-^/Atg5^fl/fl^*(n=4), or *Caspase1/11^-/-^/Atg5^fl/fl^-LysM-Cre* (n=5) mice. The legend in 4A should be used for 4B too. **(C)** Kaplan Meier curve of survival proportions during *Mtb-* GFP infection of *Atg5^fl/fl^*, *Atg5^fl/fl^-LysM-Cre*, *Caspase1/11^-/-^/Atg5^fl/fl^*, and *Caspase1/11^-/-^/Atg5^fl/fl^-LysM-Cre* mice. **(D)** Proportion of CD45^+^ cells that are neutrophils (CD45^+^Ly6G^+^CD11b^+^) in the lung at 14 dpi in *Mtb-*GFP infected wildtype (n=4), *Parkin^-/-^* (n=3) or *Pink1^-/-^* (n=3) mice reported as a ratio relative to the average neutrophil frequency in wildtype mice. **(E)** Lung burden from the right lobe of the lung at 14 dpi in *Mtb-*GFP infected wildtype (n=4), *Parkin^-/-^* (n=3) or *Pink1^-/-^* (n=3) mice. Statistical differences were determined by log-rank Mantel-Cox test (C) and one-way ANOVA and Šídák multiple comparison test (A-B, D-E) * P < 0.05, ** P < 0.01, *** P < 0.001, **** P < 0.0001. Differences that are not statistically significant are designated as ns. Pooled data from at least two separate experiments is graphed where each data point is from one biological replicate.

Autophagy has also been shown to suppress inflammatory responses via the process of mitophagy, where autophagy targets old and damaged mitochondria to the lysosome for degradation (56,57). The build-up of damaged or dysfunctional mitochondria in the absence of autophagy results in loss of mitochondrial membrane potential and the release of reactive oxygen species (ROS), mitochondrial DNA, and ATP to the cytosol where it can lead to oxidative damage, inflammasome activation, and pro-inflammatory cytokine production (57–61). Mitophagy requires the canonical autophagy proteins, including ATG5, ATG16L1, and BECLIN 1, as well as PARKIN and PTEN-induced kinase 1 (PINK1) (62). PINK1 accumulates on damaged mitochondria and activates the mitochondrial E3 ubiquitin ligase, PARKIN, to ubiquitinylate damaged mitochondria (58,61). Optineurin and NDP52 are the main mitophagy receptors that interact with the ubiquitinylated mitochondria and LC3, leading to autophagosome engulfment of the mitochondria (58,63). To investigate whether loss of mitophagy could contribute to higher neutrophil accumulation in the lungs following *M. tuberculosis* infection, we measured neutrophil frequency in the lungs at 14 dpi by flow cytometry in *Parkin^-/-^* and *Pink1^-/-^* mice relative to WT controls. There was no difference in neutrophil abundance or bacterial burdens in *M. tuberculosis*-infected *Parkin^-/-^* or *Pink1^-/-^* mice relative to WT mice at 14 dpi **(Fig 4D and 4E)**, indicating that mitophagy is not required to control neutrophil recruitment early during *M. tuberculosis* infection. To determine if mitophagy is required in macrophages to control proinflammatory cytokine and chemokine production during *M. tuberculosis* infection we generated BMDMs from *Parkin^-/-^, Pink1^-/-^*, and WT mice and infected the macrophages with *M. tuberculosis* for 24 hours. We measured cytokine and chemokine levels from mock and *M. tuberculosis* infected cultures using the cytokine bead array (Bio-Rad). Unlike in autophagy-deficient BMDMs, there were no differences in IL-6, IL-1β, G-CSF, KC, TNF-α or RANTES production by *M. tuberculosis* infected *Parkin^-/-^* and *Pink1^-/-^* macrophages at 24 hpi compared to WT macrophages (**S3 Fig**), nor any differences in bacterial burden (**S3 Fig**). Therefore, loss of mitophagy in macrophages does not result in higher levels of inflammation during *M. tuberculosis* infection.

### ATG5 is required to suppress early T_H_17 responses in the lungs during *M. tuberculosis* infection

The higher levels of IL-17A observed in the lungs of *Atg5^fl/fl^-LysM-Cre* mice at 14 dpi with *M. tuberculosis* relative to controls was not reproduced by BMDMs infected with *M. tuberculosis* for 24 hours (**S2 Fig)**. Although there are many reasons to explain this, one possibility is that the macrophages were not the source of IL-17A *in vivo*. We investigated what cell type was expressing higher levels of IL-17A in the *Atg5^fl/fl^-LysM-Cre* mice during *M. tuberculosis* infection by crossing the *Atg5^fl/fl^* and *Atg5^fl/fl^-LysM-Cre* mice with an IL-17A reporter mouse that expresses GFP under the IL-17A promoter (Jax # 018472). We infected *Il17a-GFP/Atg5^fl/fl^*and *Il17a-GFP/Atg5^fl/fl^-LysM-Cre* mice with *M. tuberculosis* and monitored GFP expression as a proxy of IL-17A expression in immune cells at 14 dpi. The only cell type we reproducibly detected >0.5% of the cells expressing GFP were CD4^+^ T cells. Similar to previous studies with *Atg5^fl/fl^* and *Atg5^fl/fl^-LysM-Cre* mice, there was no difference in total CD4^+^ T cell numbers in the lungs of *Il17a-GFP/Atg5^fl/fl^* and *Il17a-GFP/Atg5^fl/fl^-LysM-Cre* mice at 14 dpi **(Fig 5A)** (3). However, a greater frequency and number of the CD4^+^ T cells in the lungs of *Il17a-GFP/Atg5^fl/fl^-LysM-Cre* mice at 14 dpi were IL-17-GFP^+^ compared to *Il17a-GFP/Atg5^fl/fl^* mice **(Fig 5B and 5C)**. These data indicate that CD4^+^ T cells contribute to the higher levels of IL-17A in the lungs of *M. tuberculosis* infected *Atg5^fl/fl^-LysM-Cre* mice and ATG5 is required in innate immune cells to negatively regulate T_H_17 responses during *M. tuberculosis* infection.

**Figure 5.**
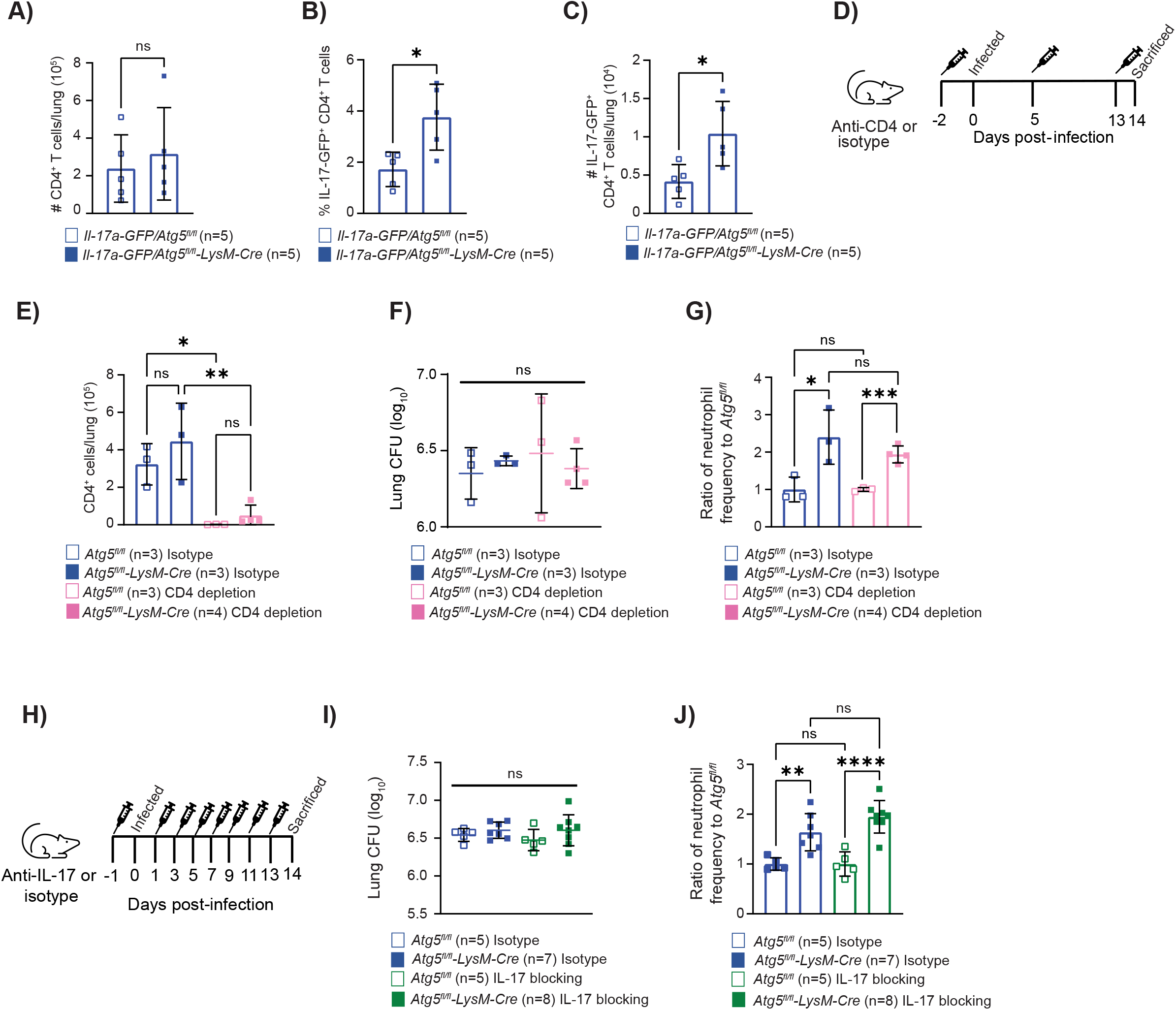
ATG5 is required to suppress early T_H_17 responses during *M. tuberculosis* infection. **(A)** The number of CD4^+^ T cells (CD45^+^ TCRβ^+^ CD4^+^) in *Atg5^fl/fl^* and *Atg5^fl/fl^-LysM-Cre* mice are reported as the total cells per lung in the left lobe at 14 dpi. **(B)** The frequency of IL-17-GFP^+^ CD4^+^ T cells in the lung (CD45^+^ TCRβ^+^ CD4^+^ IL-17-GFP^+^) of *Atg5^fl/fl^* and *Atg5^fl/fl^-LysM-Cre* mice are reported as the percentage of CD4^+^ T cells that are IL-17-GFP positive at 14 dpi. **(C)** The number IL-17-GFP^+^ CD4^+^ T cells in *Atg5^fl/fl^* and *Atg5^fl/fl^-LysM-Cre* mice are reported as the total cells per lung in the left lobe at 14 dpi. **(D)** Schematic depicting the timing of CD4-depletion antibody injections. **(E)** The number of CD4^+^ T cells (CD45^+^ TCRβ^+^ CD4^+^) in *Mtb-*GFP infected *Atg5^fl/fl^* and *Atg5^fl/fl^-LysM-Cre* mice are reported as the total cells per lung in the left lobe at 14 dpi following antibody treatment. **(F)** Lung burden from the right lobes of the lung at 14 dpi in *Mtb-*GFP infected *Atg5^fl/fl^* or *Atg5^fl/fl^-LysM-Cre* mice that received isotype or CD4-depletion antibodies. **(G)** Proportion of CD45^+^ cells that are neutrophils (CD45^+^Ly6G^+^CD11b^+^) in the lung at 14 dpi in *Mtb-*GFP infected *Atg5^fl/fl^* or *Atg5^fl/fl^-LysM-Cre* mice that received CD4-depletion or isotype treatment. Neutrophil frequency is reported as a ratio relative to the average neutrophil frequency in *Atg5^fl/fl^*control mice at 14 dpi. **(H)** Schematic depicting delivery of the IL-17 neutralizing antibody treatments. **(I)** Lung burden from the right lobes of the lung at 14 dpi in *Mtb-*GFP infected *Atg5^fl/fl^* or *Atg5^fl/fl^-LysM-Cre* mice that received isotype or IL-17 neutralizing antibodies. **(J)** Proportion of CD45^+^ cells that are neutrophils (CD45^+^Ly6G^+^CD11b^+^) in the lung at 14 dpi in *Mtb-*GFP infected *Atg5^fl/fl^* or *Atg5^fl/fl^-LysM-Cre* mice that received isotype or IL-17 neutralizing antibodies. Neutrophil frequency is reported as a ratio relative to the average neutrophil frequency in *Atg5^fl/fl^* control mice at 14 dpi. Statistical differences were determined by student t-test (A) or one-way ANOVA and Šídák multiple comparison test (A-C, E-G, I and J). * P < 0.05, ** P < 0.01, *** P < 0.001, **** P < 0.0001. Differences that are not statistically significant are designated as ns. Pooled data from at least two separate experiments is graphed where each data point is from one biological replicate.

IL-17A drives neutrophil influx by promoting the production of neutrophil chemokines MIP-1α and KC and through activation of endothelial cells (64–66). Therefore, the increased T_H_17 responses could be responsible for the early influx and accumulation of neutrophils in the lungs of *M. tuberculosis*-infected *Atg5^fl/fl^-LysM-Cre* mice. To determine if the increased expression of IL-17A by T cells was responsible for the enhanced influx of neutrophils at 14 dpi in *Atg5^fl/fl^-LysM-Cre* mice, we depleted CD4^+^ T cells by administering antibodies specific for CD4 from day -2 to 14 dpi **(Fig 5D and 5E)**. At 14 dpi, we harvested the lungs for enumeration of *M. tuberculosis* burden and neutrophil abundance and found that there was no effect of CD4^+^ T cell depletion on either readout **(Fig 5F and 5G)**. In addition, blocking IL-17A signaling by administering an anti-IL-17A antibody from day -1 to 14 dpi **(Fig 5H)** did not affect *M. tuberculosis* burden **(Fig 5I)** or neutrophil abundance **(Fig 5J)** in *Atg5^fl/fl^-LysM-Cre* mice and *Atg5^fl/fl^* mice at 14 dpi. Therefore, although ATG5 is required in innate immune cells to suppress IL-17A expression in T cells, this role does not contribute to differences in neutrophil accumulation early during *M. tuberculosis* infection.

## DISCUSSION

*Atg5^fl/fl^-LysM-Cre* mice are extremely susceptible to *M. tuberculosis* infection, where neutrophils accumulate in the lungs of infected *Atg5^fl/fl^-LysM-Cre* mice by 14 dpi and are sustained at high levels until the mice succumb to the infection between 30-40 dpi (3). It was previously unknown how ATG5 imparted control of neutrophil recruitment to the lungs during *M. tuberculosis* infection. We have discovered that ATG5 is required in CD11c^+^ lung macrophages and DCs to regulate proinflammatory cytokine production and neutrophil accumulation in the lungs during *M. tuberculosis* infection. This role for ATG5 is shared with ATG16L1 and BECLIN 1, but not RUBICON, indicating it is autophagy-dependent and does not involve LAP. *Atg5^fl/fl^-CD11c-Cre* mice succumb to *M. tuberculosis* infection at a similar time as *Atg16l1^fl/fl^-LysM-Cre* and *Atg7^fl/fl^-LysM-Cre* mice (23), suggesting that loss of the role for autophagy that we have identified in CD11c^+^ lung macrophages and DCs is responsible for the susceptibility of *Atg16l1^fl/fl^-LysM-Cre* and *Atg7^fl/fl^-LysM-Cre* mice to *M. tuberculosis*. These data demonstrate that autophagy is required in CD11c^+^ cells to control inflammatory response during *M. tuberculosis* infection and promote survival through the chronic phase of infection. We were able to reproduce the heightened proinflammatory responses in *M. tuberculosis* infected autophagy-deficient BMDMs *in vitro*, suggesting that autophagy specifically suppresses inflammatory responses from macrophages during *M. tuberculosis* infection, although this does not rule out a similar role in DCs *in vivo*. Alveolar macrophages are among the first cells to encounter *M. tuberculosis* in the airways and orchestrate the initial response to infection, recruiting other innate immune cells to the lung (35,36). We postulate that similar to the BMDMs, autophagy-deficient alveolar macrophages overproduce pro-inflammatory signals during *M. tuberculosis* infection, leading to increased neutrophil recruitment.

The increased levels of cytokines and chemokines produced by autophagy-deficient macrophages was dependent on *M. tuberculosis* infection, demonstrating that pathogen detection was required. However, the heightened proinflammatory responses in autophagy-deficient macrophages occurred in the absence of differences in bacterial burden, indicating that the enhanced inflammatory response is not due to increased antigen. Although multiple studies have shown effects of loss of autophagy on *M. tuberculosis* burden in macrophages and DCs *in vitro* in cell culture (5,23–29), there was no difference in *M. tuberculosis* burden in these cell types *in vivo* when we detected higher levels of proinflammatory cytokine production and neutrophil influx. Therefore, the role for autophagy in regulating proinflammatory cytokine production from CD11c^+^ lung macrophages and DCs during *M. tuberculosis* infection is independent of xenophagy controlling *M. tuberculosis* replication. We also ruled out the involvement of CASPASE1/11-dependent inflammasome activity and mitophagy in autophagy-dependent regulation of neutrophil accumulation during *M. tuberculosis* infection, leaving open the question of how autophagy regulates macrophage proinflammatory responses during infection.

Our understanding of the number of cellular pathways regulated by autophagy continues to grow and there are many possible mechanisms by which autophagy could regulate inflammatory responses in macrophages (18). One possibility is that targeting of autophagy components to intracellular *M. tuberculosis* prevents sensing of *M. tuberculosis* by cytosolic pattern recognition receptors and subsequent proinflammatory cytokine and chemokine production, without impacting *M. tuberculosis* replication. Autophagy can also regulate levels of IL-1β (44,62,67), IL-6 (32,68,69) and TNF-α (68,69) downstream of pathogen sensing, in part by negatively regulating NFκB activation (70–72). Autophagy can both directly promote autophagic cell death and apoptosis as well as negatively regulate RIPK3-dependent necrotic cell death, all which can impact inflammatory responses (23,73–77). Although we did not detect overall differences in the viability of autophagy deficient cells at the time points we observed effects on cytokine and chemokine production *in vivo* or *in vitro*, the mechanism of cell death in *Atg5^fl/fl^* and *Atg5^-/-^* cells during *M. tuberculosis* infection may vary and could impact the inflammatory responses (73,74,78). Autophagy may also be required in macrophages for the efficient removal of dead or infected neutrophils from the lung and resolution of inflammation. The best studied process for removal of dead cells by phagocytes is efferocytosis (79). LAP has specifically been shown to contribute to efferocytosis (80), and although we rule out a role for LAP in regulating neutrophil accumulation during *M. tuberculosis* infection, in remains possible that processing of dead cells may be attenuated in the absence of autophagy. Another possible mechanism for how autophagy regulates inflammatory responses from macrophages during *M. tuberculosis* infection involves the process of ER-phagy. ER-phagy is induced under conditions of ER stress, accumulation of unfolded proteins, and during infection (81,82). ER-phagy restrains ER stress responses by targeting excess or damaged endoplasmic reticulum to autophagosomes for degradation (81), but in the absence of autophagy, ER stress activates NFκB-dependent transcription of inflammatory cytokines, such as IL-1β, IL-6, IL-18 and TNF-α (82).

We also discovered that ATG5 was required in innate immune cells to suppress early T_H_17 responses during *M. tuberculosis* infection. At this point, we do not know if the increased abundance of IL-17^+^ CD4^+^ T cells in *M. tuberculosis* infected *Atg5^fl/fl^-LysM-Cre* mice is due to an autophagy-dependent or independent role for ATG5, but there are multiple other reports of autophagy regulating T_H_17 responses. The effect of loss of *Atg5* in innate immune cells on IL-17A expression from T cells may be explained by the requirement for autophagy in DCs to negatively regulate surface expression of disintegrin and metalloproteinase domain-containing protein 10 (ADAM10). ADAM10 cleaves its substrate ICOSL and lower levels of ICOSL leads to decreased ICOSL-ICOS interactions between DCs and T cells, resulting in less CD25^hi^ CD4^+^ T regulatory cells and more IL-17^+^ CD4^+^ T cells (83). In addition, autophagy negatively regulates T_H_17 differentiation by reducing IL-23 and IL-1β levels, which promote T_H_17 differentiation and IL-17A secretion (4,84–86). Nonetheless, depleting CD4^+^ T cells or blocking IL-17A did not rescue the increased neutrophil accumulation in the lungs of *M. tuberculosis* infected *Atg5^fl/fl^-LysM-Cre* mice at 14 dpi, suggesting that the hyper-inflammatory responses from infected autophagy-deficient macrophages and DCs is sufficient to recruit excessive neutrophils early during infection. However, it is possible that the heightened T_H_17 responses in *M. tuberculosis* infected *Atg5^fl/fl^-LysM-Cre* mice have a longer-term impact on the increased susceptibility of these mice and could contribute to the susceptibility of *Atg5^fl/fl^-Cd11c-Cre* mice during the chronic phase of infection.

Our data support a role for autophagy in restricting neutrophil accumulation in the lung but does not rule out additional contributions from autophagy independent roles for ATG5, BECLIN 1 and ATG16L1. In particular, loss of *Atg5, Becn1* or *Atg16l1* in LysM^+^ cells led to different degrees of elevated neutrophil frequency in the lung at 14 dpi relative to control mice, where loss of *Atg5* results in the greatest increase in neutrophil abundance. This could in part be due to differences in efficiency of gene deletion in the different lines (23). In addition, we have previously identified an autophagy independent role for ATG5 in neutrophils, where loss of ATG5 in neutrophils can result in early lethality during *M. tuberculosis* infection (3). Therefore, loss of *Atg5* in LysM^+^ cells, which includes neutrophils, may result in higher neutrophil frequency in the lung at 14 dpi than loss of *Becn1* or *Atg16l1* because of this autophagy-independent role for ATG5 in neutrophils. We hypothesize that the combination of the newly discovered role for autophagy in CD11c^+^ lung macrophages and DCs to regulate inflammatory responses and an autophagy-independent role for ATG5 in neutrophils collectively allow for control of *M. tuberculosis* infection, where loss of both functions results in the extreme susceptibility of *Atg5^fl/fl^-LysM-Cre* mice to *M. tuberculosis* infection. Higher abundances of neutrophils have been associated with poor disease prognosis and treatment outcomes in TB patients (14–16). Therefore, our new findings and future dissection of the ATG5-dependent mechanisms of regulating neutrophil recruitment to the lungs during *M. tuberculosis* infection will provide critical insight into how to promote protective immune responses to TB.

## MATERIALS AND METHODS

### Mice

All flox mice (*Atg5^fl/lfl^*, *Atg16l1^fl/fl^*, *Becn1^fl/fl^*) used in this study have been described previously (3,87–89) and colonies are maintained in an enhanced barrier facility. LysM-Cre (Jax #004781), Cd11c-Cre (Jax #007567), Mrp8-Cre (Jax #021614) from the Jackson Laboratory were crossed to specific flox mice. *Il17a-IRES-GFP-KI* (Jax # 018472) reporter mice were bred to *Atg5^fl/fl^-LysM-Cre* and *Atg5^fl/fl^* mice to generate the *Il17a-GFP/Atg5^fl/fl^-LysM-Cre* and *Il17a-GFP/Atg5^fl/fl^* lines. *Rubicon^-/-^*(Jax # 032581) and WT litter mates were provided by D. Green and J. Martinez (90). Caspase 1/11^-/-^ (Jax #016621) were bred to *Atg5^fl/fl^-LysM-Cre* and *Becn1^fl/fl^-LysM-Cre* mice. *Parkin^-/-^* (Jax # 006582)(91), *Pink1^-/-^* (Jax # 017946) (56), and WT control mice were a gift from Dr. Jonathan Brestoff at Washington University School of Medicine. Male and female littermates (aged 6-12 weeks) were used and were subject to randomization. A minimum of 3 mice were used per experiment and each experiment was performed twice. Statistical consideration was not used to determine mouse sample sizes. The mice were housed and bred at Washington University in St. Louis in specific pathogen-free conditions in accordance with federal and university guidelines, and protocols were approved by the Animal Studies Committee of Washington University.

### Infection of mice with *M. tuberculosis* and measurement of bacterial burden in the lungs

*M. tuberculosis* Erdman expressing GFP (*Mtb*-GFP (10,92)) was used in all experiments except experiments with the *Il-17a-GFP*/*Atg5^fl/fl^-LysM-Cre* reporter mice when WT Erdman was used. *M. tuberculosis* was cultured at 37°C in 7H9 (broth) or 7H11 (agar) (Difco) medium supplemented with 10% oleic acid/albumin/dextrose/catalase (OADC), 0.5% glycerol, and 0.05% Tween 80 (broth). Cultures of GFP expressing *M. tuberculosis* were grown in the presence of kanamycin (20μg/mL) to ensure plasmid retention. *M. tuberculosis* cultures in logarithmic growth phase (OD600 = 0.5–0.8) were washed with PBS + 0.05% Tween-80, sonicated to disperse clumps, and diluted in sterile water before delivering 100 CFUs of aerosolized *M. tuberculosis* per lung using an Inhalation Exposure System (Glas-Col). Within 2 hours of each infection, lungs were harvested from at least two control mice, homogenized, and plated on 7H11 agar to determine the input CFU dose. At 14 dpi, *M. tuberculosis* titers were determined by homogenizing the superior, middle, and inferior lobes of the right lung and plating serial dilutions on 7H11 agar. Colonies were counted after 3 weeks of incubation at 37°C in 5% CO2.

### Flow cytometry from blood and infected lungs

Blood was collected by cardiac puncture into K2EDTA anticoagulant tubes (BD, 365974). Red blood cell lysis was performed by adding 2ml of ACK lysis buffer (Gibco, A10492-01) per 100µL of EDTA-treated blood for 5 minutes at room temperature. Cells were pelleted at 500xg for 5 minutes and then resuspended in 50uL of PBS+ 2% HI-FBS + 2mM EDTA in the presence of Fc receptor blocking antibody (BioLegend, 101302). Cells were labelled with antibodies as described below. Lungs were perfused with sterile PBS and digested for 1 hour with 625µg/mL collagenase D (Roche 11088875103) and 75U/mL DNase I (Sigma D4527). Cells were quenched with PBS + 2% heat-inactivated (HI)-FBS, + 2mM EDTA and passed through a 70µM filter. Cells were stained with Zombie-violet or Zombie-NIR in PBS at 1:2000 dilution for 5 minutes at room temperature prior to resuspending in PBS + 2% HI-FBS + 2mM EDTA in the presence of Fc receptor blocking antibody (BioLegend, 101302) for blocking. Cells were labelled with antibodies at a 1:200 dilution using the following mouse markers: CD11b_BV605 or PerCP-Cy5.5 (clone M1/70), CD45_AF700 (BioLegend, 103259), Ly6G_PE-Cy7 or AF647 (clone 1A8), MHCII_Spark blue 550 (BioLegend, 107662), CD62L_Pe/Cy5 (BioLegend, 104410), CD44_BV510 (BioLegend, 103044), CD11c_PerCP (BioLegend, 117325), Ly6C_BV605 (BioLegend, 128036), CD4_BV570 (clone RM4-5), TCRb_BV421 (clone H57-597), CD19_Pacific blue (BioLegend, 115526), Zombie-violet or -NIR (BioLegend, 423113 or 423105), SiglecF_BV480 (BD Biosciences, 746668), MHC-II_Sparkblue550 (BioLegend, 107662), CD64_PerCP-eFluor 710 (eBiosciences, 46061482) and MerTK_PE/Cy7 (eBiosciences, 25575182). Cells were incubated for 20 minutes at 4°C with antibodies, washed and then fixed in 4% paraformaldehyde (Electron Microscope Sciences) for 20 minutes at room temperature. Flow cytometry was performed on an LSR-Fortessa (BD Bioscience) or an Aurora (Cytek Biosciences, with 4 laser 16V-14B-10YG-8R configuration) and analyzed using FlowJo software (Tree Star). Absolute cell counts were determined using Precision count beads (BioLegend) or volumetric-based counting on the Aurora. Gating strategies to identify *Mtb-*GFP*^+^*myeloid cell populations, viable lung neutrophils, CD4^+^ T cells and IL-17-GFP^+^ CD4^+^ T cells are in supplementary figure S4A-S4C.

### Culturing and infection of bone marrow derived macrophages (BMDMs)

BMDMs were generated by flushing femurs and tibias of mice and culturing the cells in DMEM, 20% HI-FBS, 10% supernatant from 3T3 cells overexpressing M-CSF + 1% MEM non-essential amino acids (Cellgro 25-025-CI), 2mM L-glutamine, 100U/mL penicillin and 100µg/mL streptomycin (Sigma P4333) at 37°C in 5% CO_2_ in non-TC treated plates. After 6 days non-adherent cells were removed and 1×10^6^ adherent macrophages were seeded per well in 6 well non-TC treated plates in DMEM, 10% HI-FBS, 1% MEM non-essential amino acids and 2 mM L-glutamine. BMDMs were rested overnight at 37°C in 5% CO_2_. *M. tuberculosis* was grown to an OD of 0.6-0.8, washed with PBS twice, sonicated to disperse clumps, centrifuged at 55xg for 10 minutes to remove clumps and resuspended in antibiotic-free BMDM media. Macrophages were infected at an MOI of 10 by centrifuging the cells at 200xg for 10 minutes. BMDMs were washed with PBS twice to remove free *M. tuberculosis* and fresh BMDM media was added to each well. Cells were incubated at 37°C and 5% CO_2_ for 24 hours. To determine CFU counts, the cells were lysed with 0.05% triton X-100, serially diluted, and plated onto 7H11 agar and incubated for 21 days when bacterial colonies were counted. At 24 hours post-infection (hpi) supernatants were stored at -80°C for cytokine analysis. To assess BMDM viability, cells were harvested at 24 hpi by gently scrapping, washed with PBS twice, stained with Zombie-violet in PBS at 1:2000 dilution for 5 minutes at room temperature, washed once with PBS, and fixed in 4% PFA for 20 minutes at room temperature before performing flow cytometry. Flow cytometry was performed on an LSR-Fortessa (BD Bioscience) and analyzed using FlowJo software (Tree Star). The gating strategy to identify viable BMDMs at 24 hpi is depicted in Supplementary Figure 1D.

### Cytokine analysis

BMDM supernatants were filtered through a 0.22 µm filter twice to remove *M. tuberculosis* and analyzed using the BioPlex-Pro Mouse Cytokine 23-Plex Immunoassay (Bio-Rad) as per the manufacturer’s instructions. ELISAs were performed according to the manufacturer’s instructions (R&D systems): KC/CXCL1 (DY453), IL-6 (DY406) and G-CSF (DY414).

### IL-17A blocking and T cell depletion

To neutralize IL-17A, 100µg of InVivo monoclonal anti-IL-17A (Bio X Cell, BE0173) neutralizing antibody was administered to *Atg5^fl/fl^* and *Atg5^fl/fl^-LysM-Cre* mice by intraperitoneal (i.p.) injection every other day starting at 1 day prior to infection, with the final dose delivered at 13 dpi, similar to published protocols (93). Control mice received 100µg of IgG from mouse serum (Sigma, I5381) by i.p. injection every other day starting 1 day prior to infection and finishing on 13 dpi. To deplete CD4^+^ T cells from mice, 250µg of anti-mouse CD4 (Leinco Technologies, C1333) was administered by i.p. injection at 2 days prior to infection, 5 dpi and 12 dpi. Control mice received 250µg of IgG from rat serum (Sigma, 18015) i.p. on 2 days prior to infection, 5 dpi and 12 dpi.

### Data and statistics

All experiments were performed at least twice. When shown, multiple samples represent biological (not technical) replicates of mice randomly sorted into each experimental group. No blinding was performed during animal experiments. Animals were only excluded when pathology unrelated to *M. tuberculosis* infection was present (i.e. bad teeth leading to weight loss). Determination of statistical differences was performed with Prism (GraphPad Software, Inc.) using log-rank Mantel-Cox test (survival), unpaired two-tailed t-test (to compare two groups with similar variances), or one-way ANOVA with Šídák Multiple Comparison test (to compare more than two groups). When used, center values and error bars represent the mean +/- S.E.M. In all figures, all significant differences are indicated by asterisks: * *P<0.05, ** P<0.01, *** P<0.001, **** P<0.0001.* Non-significant comparisons of particular interest are noted as n.s.

## ACKNOWLEDGEMENTS

The authors thank Dr. Jonathan Brestoff at Washington University School of Medicine for generously providing the *Parkin^-/-^* and *Pink1^-/-^* mice. This work was supported by NIH grants R01 AI132697 and U19 AI142784, a Burroughs Wellcome Fund Investigators in the Pathogenesis of Infectious Disease Award, and the Philip and Sima Needleman Center for Autophagy Therapeutics and Research to C.L.S., a Potts Memorial Foundation postdoctoral fellowship to R.L.K., and a National Science Foundation Graduate Research Fellowship DGE-1143954 and the NIGMS Cell and Molecular Biology Training Grant GM007067 to J.M.K.

## AUTHOR CONTRIBUTIONS

The experiments were designed by R.L.K., J.M.K., and C.L.S. The experiments were executed by R.L.K., J.M.K., and A.S. with assistance from R.W., M.R.G. and S.M.C. D.K. bred and maintained the mouse colonies. The manuscript was written by R.L.K. and C.L.S.

## FIGURE LEGENDS

**Supplemental Figure 1. ATG5 is not required in CD11c^+^ cells to regulate neutrophil viability in the lung or accumulation of neutrophils in the blood during *M. tuberculosis* infection. (A)** The proportion of lung neutrophils (CD45^+^ Ly6G^+^ CD11b^+^) that are viable (Zombie^-^) in *M. tuberculosis* infected *Atg5^fl/fl^* (n=8) and *Atg5^fl/fl^-CD11c-Cre* (n=7) mice at 14 dpi. **(B)** The proportion of CD45^+^ cells in the blood that are neutrophils in *Atg5^fl/fl^*(n=4 in naïve and n=3 in 14 dpi) and *Atg5^fl/fl^-CD11c-Cre* (n=4) mice at 14 dpi and uninfected (naïve) mice. Statistical differences were determined by student t-test comparing the genotypes within a particular cell type or treatment group. * P < 0.05, ** P < 0.01, *** P < 0.001, **** P < 0.0001. Differences that are not statistically significant are designated as ns. Pooled data from at least two separate experiments are graphed where each data point is from one biological replicate.

**Supplemental Figure 2. Levels of viability, cytokines, and chemokines in autophagy-deficient macrophages during *M. tuberculosis* infection. (A)** The proportion of *Atg5^fl/fl^* (n=5) and *Atg5^fl/fl^-LysM-Cre* (n=5) BMDMs that are viable (Zombie^-^) at 24 hpi in mock and *Mtb-*GFP treated groups. Cytokine bead array data from mock treated and *Mtb-*GFP infected BMDMs. *Atg5^fl/fl^, Atg5^fl/fl^-LysM-Cre, Atg16l1^fl/fl^, Atg16l1^fl/fl^-LysM-Cre, Becn1^fl/fl^* and *Becn1^fl/fl^-LysM-Cre* BMDMs were cultured for 24 hpi and cytokine levels were measured in the spent media from mock treated or infected macrophages. BMDMs from at least 3 mice were tested in duplicate to quantify the cytokines in the bead array. All cytokine and chemokine data that are not significantly different between *Atg5^fl/fl^-LysM-Cre* and *Atg5^fl/fl^* mice are reported here. **(B)** IL-1α, **(C)** IL-4, **(D)** IL-10, **(E)** IL-12(p40), **(F)** IL-12(p70), **(G)** IFN-γ, **(H)** MIP1β, G**(I)** M-CSF, M**(J)** IP1α, **(K)** MCP-1, **(L)** Eotaxin and **(M)** IL-17 levels at 24 hpi. Statistical differences were determined by one-way ANOVA and Šídák multiple comparison test (A) and student t-test comparing the autophagy deficient macrophage with its floxed control within a treatment condition (B-M). * P < 0.05, ** P < 0.01, *** P < 0.001, **** P < 0.0001. Cytokine levels below detection limits are designated as dbl. Differences that are not statistically significant are designated as ns. Each data point is one biological replicate, and the samples were generated from at least two separate experiments.

**Supplemental Figure 3. Mitophagy is not required in macrophages to regulate proinflammatory responses during *M. tuberculosis* infection.** WT*, Parkin^-/-^* and *Pink1^-/-^,* BMDMs were cultured for 24 hpi and cytokine levels were measured by cytokine bead array in the media from mock treated or *Mtb-* GFP infected macrophages. **(A)** IL-1β, **(B)** G-CSF, **(C)** IL-6, **(D)** KC, **(E)** RANTES and **(F)** TNF-α levels at 24 hpi **(G)** BMDM CFU counts from 24 hpi. BMDMs from at least 3 mice were tested in duplicate to quantify cytokines in the bead array. Each point is one biological replicate. Statistical differences were determined by one-way ANOVA and Šídák multiple comparison test (A-G). * P < 0.05, ** P < 0.01, *** P < 0.001, **** P < 0.0001. Cytokine levels below detection limits are designated as dbl. Statistical differences that are not significant are designated as ns. Each data point is one biological replicate, and the samples were generated from at least two separate experiments.

**Supplemental Figure 4. Gating strategies for flow cytometry identification of myeloid cells and IL-17 expressing T cells in the lung or viable BMDMs.** Representative flow cytometry plots depicting the gating strategy used to identify *Mtb-*GFP^+^ myeloid cells **(A)**, viable lung neutrophils **(B)** and IL-17-GFP^+^ CD4^+^ T cells **(C)** in the lung at 14 dpi. **(D)** Representative flow cytometry plots showing the gating strategy to identify viable BMDMs at 24 hpi. FSC-A, forward scatter area; SSC-A, side scatter area; FSC-H, forward scatter height; FSC-W, forward scatter width, Alv. Mac., alveolar macrophages; Eos., eosinophils; DC, dendritic cell, and Inf. Mono., inflammatory monocytes.

