## Supplementary figures and images for "Autophagy is required to prevent early pro-inflammatory responses and neutrophil recruitment during *Mycobacterium tuberculosis* infection without affecting pathogen replication in macrophages"

### Supplemental Figure 1

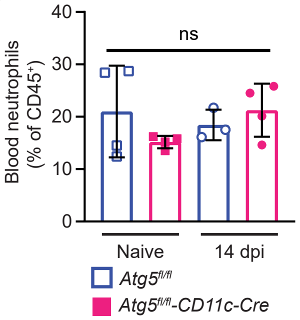

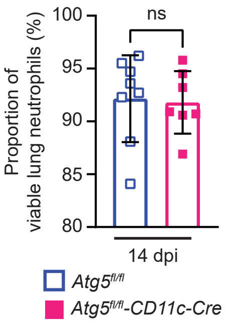

### Supplemental Figure 2

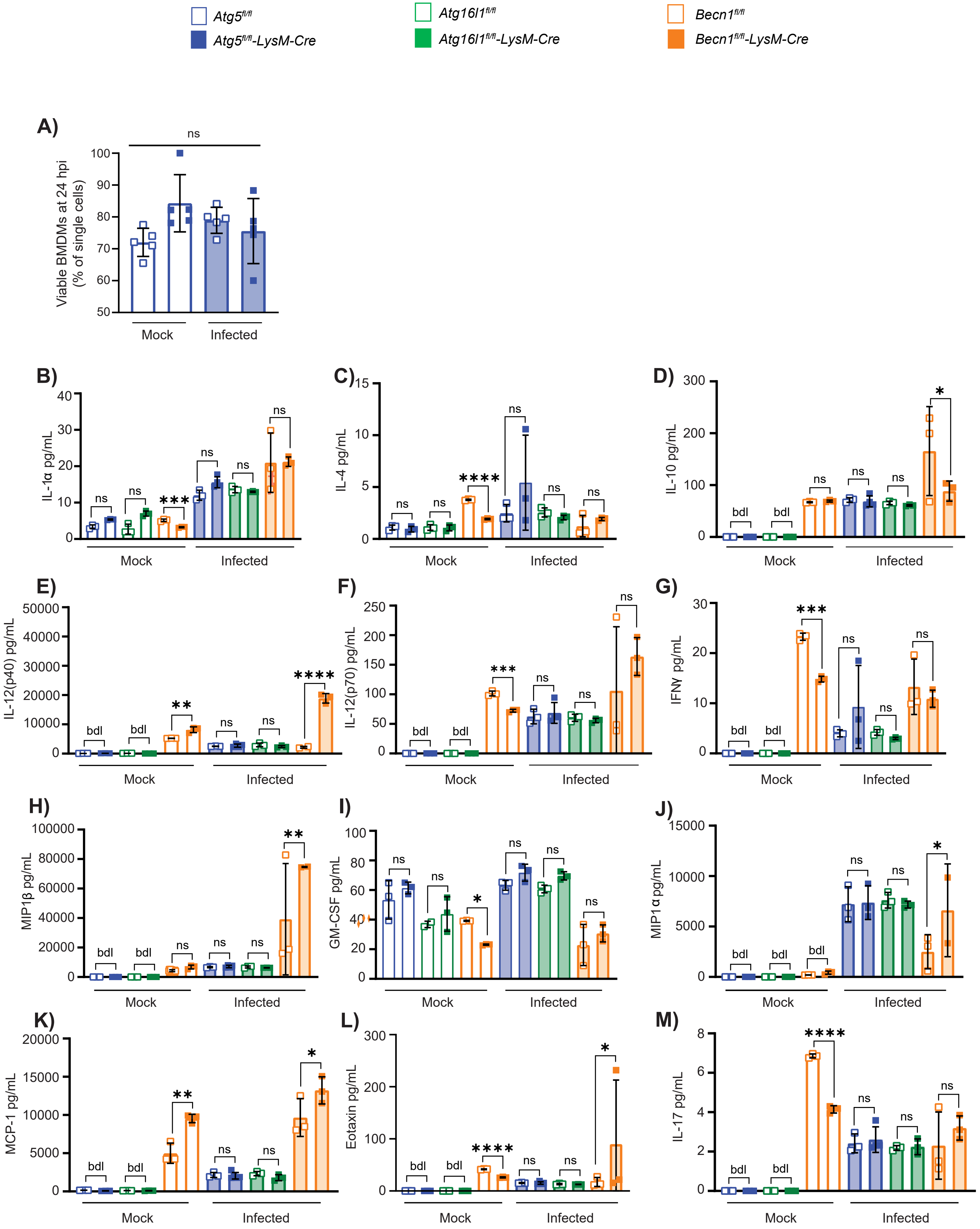

### Supplemental Figure 3

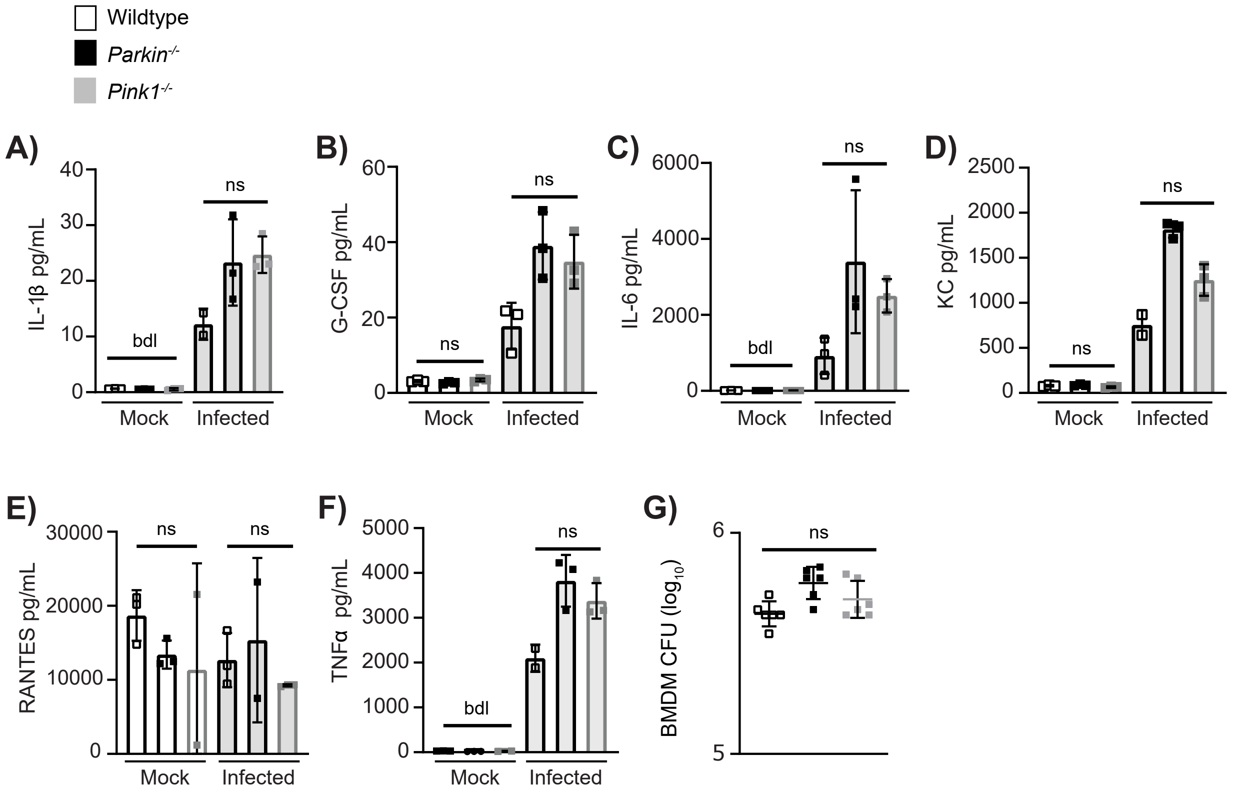

### Supplemental Figure 4

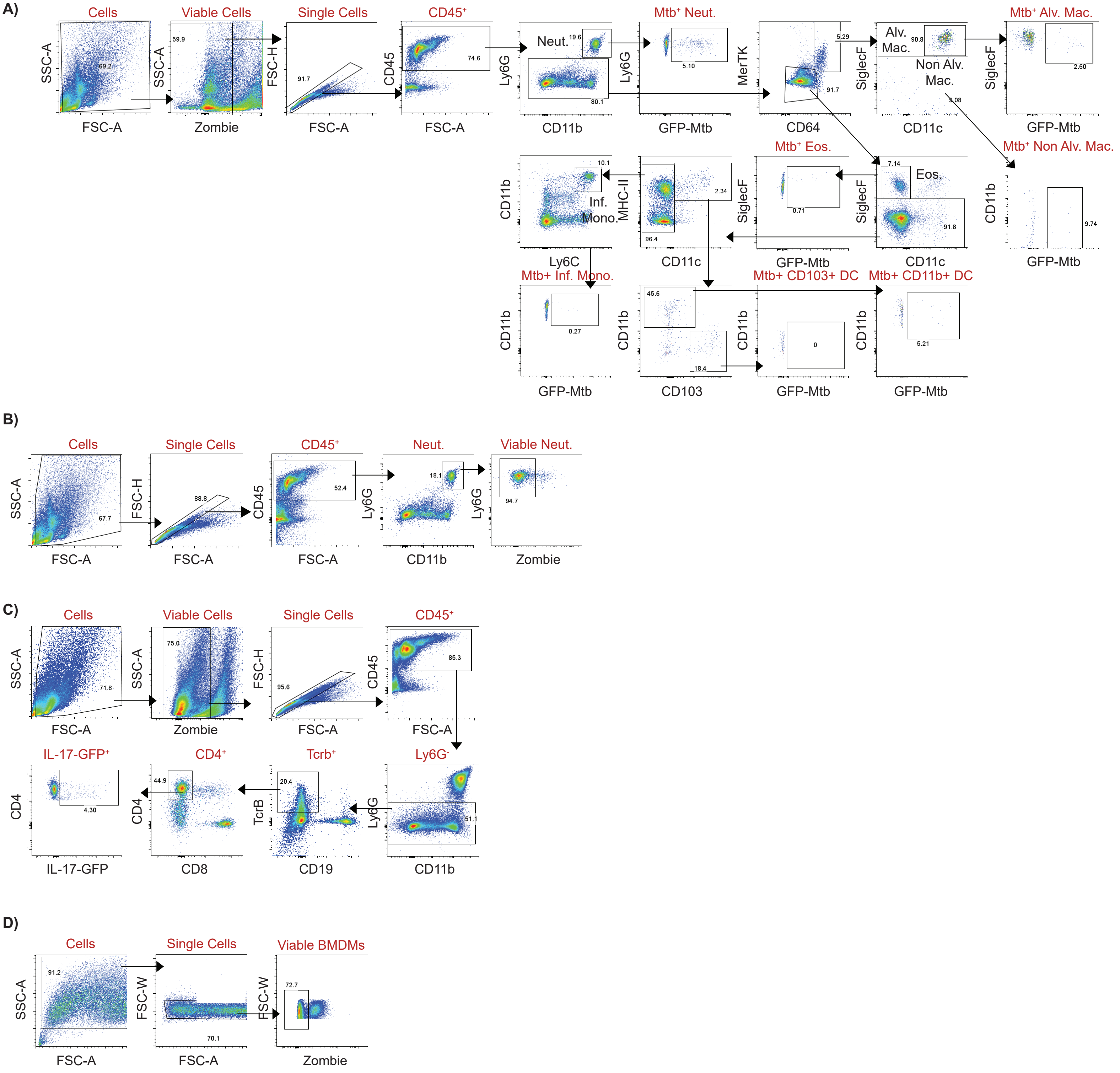
